# When migration leaves a clean trace: Decoupling migration from coalescence in the structured serial coalescent

**DOI:** 10.1101/2025.10.10.681523

**Authors:** Hao Shen, John Novembre

## Abstract

With the rapid accumulation of population genomic data across space and time, there is an urgent need for demographic inference methods that incorporate explicit time-series modeling, achieve high spatial scalability, and ensure clear identifiability between migration and coalescence rates. To address this need, we investigate pairwise genealogical processes under the structured serial coalescent, deriving evolution equations for pairwise branch length distributions and related statistics. By classifying the resulting identities according to their parameter dependencies and computational complexity, we identify a class that is not only computationally tractable but also determined exclusively by migration rates. Building on this theoretical basis, we propose a scalable framework for inferring time-varying migration rates and demonstrate its feasibility through simulation. We further outline how this framework can be extended to the joint estimation of migration and coalescence rates.

## Introduction

Since Kingman introduced the coalescent (Kingman, 1982a,b), it has been widely used in population genetic theory and inference. However, the standard coalescent theory typically assume contemporaneous sampling and panmixia, whereas empirical samples can come from different time periods and geographically structured populations with variable lineage migration and coalescence rates. These limitations have motivated key extensions: the serial coalescent for temporal stratification (Rodrigo and Felsenstein, 1999) and the structured coalescent for population subdivision (Notohara, 1990; Takahata, 1991). For models combining both temporal and spatial structure, terminology varies in the literature. Some authors retain “structured coalescent” (Müller et al., 2017) while others use “structured serial coalescent” (Ewing and Rodrigo, 2007). For clarity and distinction, we adopt the latter term throughout this paper.

These extensions of the standard coalescent have enabled powerful demographic inference tools, particularly for estimating coalescent effective population sizes (e.g., the inverse of the instantaneous local coalescence rates) and migration rates. For the serial coalescent, approaches based on full gene genealogies using tools from Bayesian phylogenetics have been developed to estimate effective population sizes from temporally sampled data (Drummond et al., 2002, 2005). For the structured coalescent and structured serial coalescent, related methods using the full gene genealogy have been proposed to jointly infer migration rates and effective population sizes (Beerli and Felsenstein, 2001; Ewing et al., 2004; Volz, 2012; Vaughan et al., 2014; De Maio et al., 2015; Müller et al., 2017, 2018).

However, because these methods rely on information from the full genealogy, their computational scalability with respect to the number of lineages and demes remains limited—even with modern algorithmic advances. Consequently, demographic inference for systems with hundreds or thousands of demes remains impractical under these frameworks.

One effective way to resolve the scalability bottleneck is to use pairwise coalescence information instead of the whole genealogy. In the context of the structured coalescent, this means developing methods based on pairwise coalescence times. The distribution of pairwise coalescence times depends on both the migration rates and coalescence rates, while also being directly connected to empirical data through pairwise summary statistics. This makes pairwise coalescence times an attractive basis for demographic inference.

For example, the methods EEMS (Petkova et al., 2016), FEEMS (Marcus et al., 2021), and FRAME (Shen and Novembre, 2025) all exploit the connection between expected pairwise coalescence times and the sample covariance matrix (McVean, 2009). Meanwhile, the method MAPS (Al-Asadi et al., 2019) uses the fact that the length distribution of long pairwise shared coalescent segments (LPSC segments, also known as identity-by-descent or IBD tracts) is determined by the distribution of pairwise coalescence times (Palamara et al., 2012; Carmi et al., 2013). All of these methods are scalable to hundreds of demes. While earlier methods like EEMS, FEEMS, and MAPS relied heavily on the assumption of symmetric migration, work by Lundgren and Ralph (2019) and by Shen and Novembre (2025) with the FRAME method now enables scalable inference of asymmetric gene flow.

Despite these methodological advances, limitations persist. First, research across many domains requires time-series modeling to accommodate serial sampling and time-varying migration rates. In conservation genetics, samples collected at multiple time points can reveal how gene flow patterns change through time. In epidemiology, serially sampled viral genomes can be used to reconstruct transmission dynamics. In human evolutionary biology, ancient DNA provides genetic snapshots from different epochs, enabling reconstruction of migration histories within each period. FRAME, however, assumes migration–drift equilibrium and therefore cannot directly accommodate serial sampling or time-varying parameters. While it is possible to analyze data by dividing it into temporal slices and fitting each slice with an equilibrium model (as demonstrated in the original MAPS and FRAME papers), it remains suboptimal compared to direct modeling under the structured serial coalescent framework. Second, these methods can have difficulty distinguishing the effects of migration rates and coalescence rates, leading to identifiability issues (MAPS is an exception, but its effectiveness under asymmetric migration scenarios remains uncertain).

These limitations motivate the need for a new demographic inference method that can: (1) preserve the scalability advantages of pairwise coalescence-based approaches, (2) directly incorporate serial sampling and time-varying parameters under the structured serial coalescent framework, and (3) effectively decouple the inference of migration rates from that of coalescence rates. In this paper, we demonstrate that such a method is theoretically achievable.

A key pairwise quantity in the structured serial coalescent is the pairwise branch length, defined as the summed branch lengths from two sampled lineages to their most recent common ancestor. Unlike the structured coalescent, where the foundational equations for expected pairwise coalescence times have been well established (even before the theory’s formal development Strobeck, 1987), the analogous equations for pairwise branch lengths in the structured serial coalescent remain largely undeveloped. To bridge this theoretical gap and facilitate new inference methods, we develop the theory for pairwise branch length dynamics under the structured serial coalescent framework.

Specifically, we derive evolution equations governing the probability density functions (PDFs) of pairwise branch lengths. By solving these equations, we uncover fundamental identities among the PDFs and systematically classify them into distinct classes based on their parameter dependencies and computational complexity. This classification framework extends naturally to other quantities derived from pairwise branch lengths, as we demonstrate through parallel analysis of mean pairwise branch length dynamics and length distribution dynamics of LPSC segments.

The general applicability of this classification enables evaluation of each class’s potential utility in demographic inference. Crucially, we identify a promising class of identities that exhibits exclusive dependence on lineage migration rates and scales computationally as *O*(*d*^3^), where *d* represents the number of demes. These properties directly address the fundamental challenges identified earlier—they resolve the migration-coalescence identifiability problem while providing the high spatial scalability required for complex population structure analysis.

This theoretical breakthrough establishes a foundation for two inference strategies. First, for studies focusing specifically on gene flow dynamics, these identities enable highly scalable migration rate estimation that is decoupled from coalescence parameters. Second, joint estimation of migration and coalescence rates may also benefit from these identities. A two-step approach—first estimating migration rates using these classes, then inferring coalescence rates conditioned on the migration parameters—could improve both parameter identifiability and computational scalability, though the latter gain may be less pronounced than in migration-only inference. We outline inference frameworks for both approaches through a proof-of-concept example and demonstrate the feasibility of the migration-only approach through preliminary simulation studies.

### The model

The structured serial coalescent process models the combined effects of backward migration between demes and coalescence events occurring when lineages reside in the same deme. We represent backward migration as a continuous-time jump process on a weighted directed graph with nodes {1, 2, …, *d*}, each corresponding to a deme. The weight on the edge from node *i* to node *j* is denoted *m*_*ij*_(*t*), representing the backward migration rate from *i* to *j* at time *t*. Biologically, *m*_*ij*_(*t*) is the rate at which deme *i* receives ancestry from deme *j*. The transition rate matrix of this jump process is *Q*(*t*), and the associated Laplacian is defined as *L*(*t*) = −*Q*(*t*). The migration rates are encoded in the weighted adjacency matrix *M* (*t*) = *Q*(*t*) − diag *Q*(*t*). The coalescence rate in deme *i* is denoted by *γ*_*i*_(*t*), and all coalescence rates are collected into the vector *γ*(*t*) = (*γ*_*i*_(*t*)).

In our model, we study the pairwise genealogical process using the pairwise branch length, defined as the sum of the branch lengths of the two lineages up to their MRCA. This measure generalizes pairwise coalescence time by explicitly accounting for temporal offsets between samples and can be viewed as the distance of the two lineages in terms of branch length. For an illustration of how pairwise branch length is computed, see Fig. 1.

**Fig. 1.**
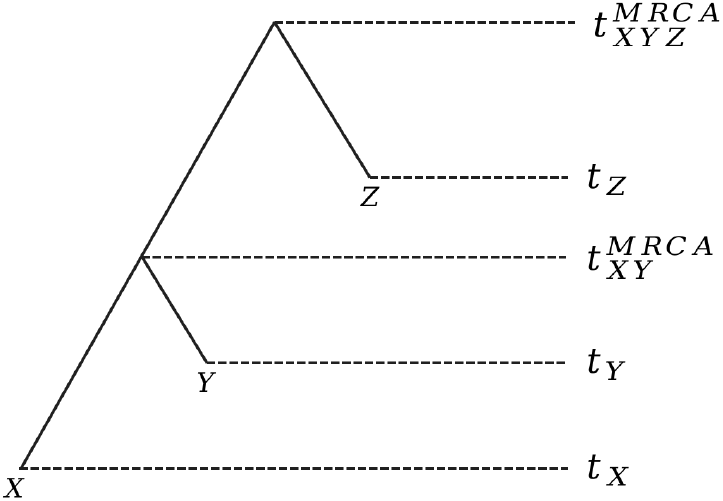
Pairwise branch length. A toy example showing how pairwise branch lengths are computed. Assume we have three samples *X, Y*, and *Z* sampled at times *t*_*X*_, *t*_*Y*_, and *t*_*Z*_, respectively. The pairwise branch length between *X* and *Y* is 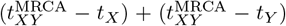. Similarly, the pairwise branch length between *X* and *Z* is 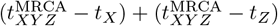, and between *Y* and *Z* is 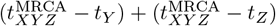.

Now let 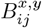 denote the random variable representing the pairwise branch length between two lineages, one sampled from deme *i* at time *x* and the other from deme *j* at time *y*, where *x* ≤ *y* and time is measured backward from the present (*t* = 0). Let 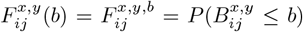 be the corresponding cumulative distribution function 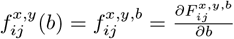 be its probability density function; 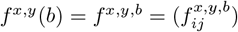 be the matrix of the probability density functions (sometimes we will neglect *b* and just write *f* ^*x,y*^ to refer to the function). All the above functions have support in [*y*− *x*, +∞) and will be 0 for *b < y* −*x*.

In the next three sections, we will explore the identities relating the probability density functions by solving the governing evolution equations, and then extend the analysis to two derived statistics (expected pairwise branch length and survival function of LPSC segment length).

### Identities relating the probability density functions of pairwise branch length

We consider a toy example with three epochs: [*t*_0_, *t*_1_), [*t*_1_, *t*_2_), and [*t*_2_, ∞), which we label as epoch 0, epoch 1, and epoch 2, respectively. The migration and coalescence rates are assumed to be piecewise constant in the first two epochs, that is, *L*(*x*) = *L*^0^, *γ*(*x*) = *γ*^0^ for *x* ∈ [*t*_0_, *t*_1_) and *L*(*x*) = *L*^1^, *γ*(*x*) = *γ*^1^ for *x* ∈ [*t*_1_, *t*_2_). We now investigate the identities relating the following time-ordered pairs of pairwise branch length PDF matrices: 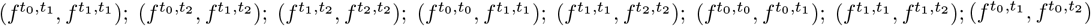. By abuse of notation, we will also use these pairs to denote the corresponding identities. These 8 identities correspond to the 8 edges in Fig. 2.

**Fig. 2.**
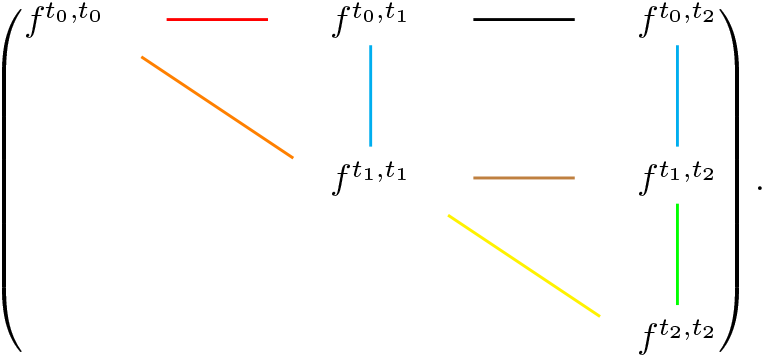
Graphical representation of the identities relating pairwise branch length probability density functions.

Graphically, these 8 edges in Fig. 2 fall naturally into three categories: the vertical edges, the diagonal edges, and the horizontal edges. In what follows, we show that this graphical classification corresponds to a classification of the identities themselves in terms of representational and computational complexity.

We first examine 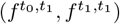, which is represented by a vertical edge. To this end, we study the evolution of 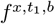 for *x* ∈ [*t*_0_, *t*_1_). As we move from *x* to *x* + Δ*t*, coalescence does not occur, and only migration of the first lineage is relevant. This yields the following equation:

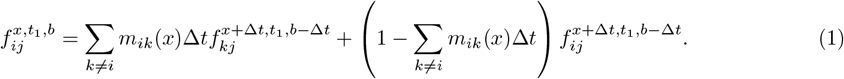

Expanding 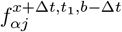 as 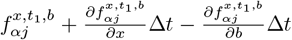 and ignoring higher-order terms, we cancel Δ*t* on both sides and obtain:

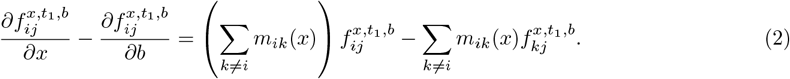

In matrix form, this partial differential equation becomes:

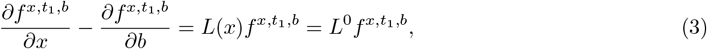

with the boundary condition given by 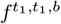. This PDE has the explicit solution:

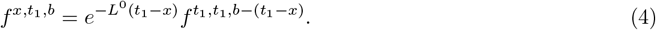

Taking the limit as *x* →*t*_0_, we obtain 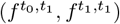. Following similar logic, 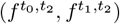 and 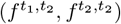, which are likewise represented by the vertical edges, can also be computed.

These three identities can be jointly expressed as:

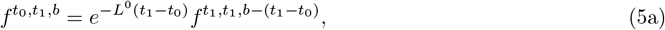

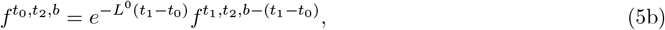

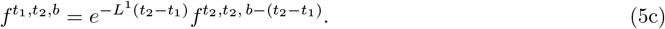

These identities are particularly valuable for two reasons. First, they depend only on backward migration rates—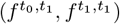 and 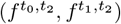 depend on *L*^0^ (we represent the corresponding edges in blue in Fig. 2), while 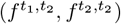 depends on *L*^1^. This makes them a promising tool for disentangling migration dynamics from coalescence effects (or effective population sizes) in complex migration–drift scenarios. Second, the computational complexity of these identities is at most *O*(*d*^3^), and can often be reduced further by exploiting the sparse structure of the Laplacian matrices. This scalability makes it possible to perform gene flow or migration inference involving hundreds of demes. As we shall see, the class of identities represented by the vertical edges is the only one that possesses these favorable properties.

Now we turn to the identity 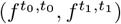, which corresponds to a diagonal edge in Fig. 2. Let *f* ^*x,b*^ = *f* ^*x,x,b*^ denote the density matrix when the two lineages are sampled at the same time. For *x* ∈ [*t*_0_, *t*_1_), the associated partial differential equation is:

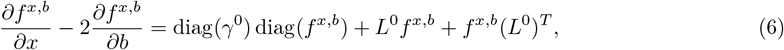

with boundary condition given by 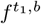 and *f* ^*x*,0^ = diag(*γ*^0^) for *x* ∈ [*t*_0_, *t*_1_). This equation is derived in a manner similar to Equation (3), with the key difference being that both lineages can migrate independently, and coalescence may occur when they are in the same deme. The PDE remains linear once we vectorize both sides.

Let *E*_*i*_ denote the matrix with a 1 in the (*i, i*)-th entry and zeros elsewhere, and let *ϵ*_*i*_ = vec(*E*_*i*_) be its vectorization. Define 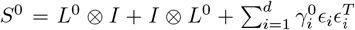, where ⊗ denotes the Kronecker product, and define 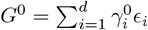. Then the solution to Equation (6) is given by:

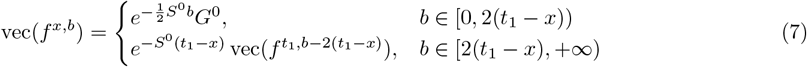

Taking the limit as *x* → *t*_0_, we obtain 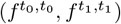. A similar analysis applies to 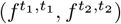, which is represented by another diagonal edge. Together, the identities represented by the diagonal edges can be written jointly as:

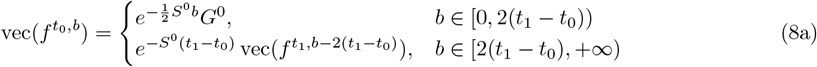

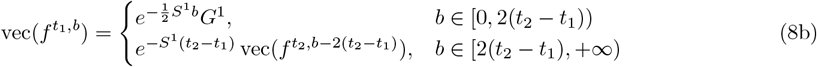

where 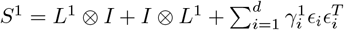 and 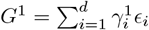.

Before turning to the identities represented by the horizontal edges, we would like to note two points.

First, when the two lineages evolve together in *f* ^*x,x,b*^, the model simplifies to a structured coalescent model. The probability density function of the pairwise coalescence times can be derived using the fact that the pairwise branch lengths are twice the pairwise coalescence times in the structured coalescent. Second, as *t*_2_ goes to infinity in Equation (8b), we obtain the stationary distribution of pairwise branch length under a structured coalescent model with migration rates coded by *L*^1^ and coalescence rates coded by *γ*^1^. The stationary distribution of pairwise branch length distribution satisfies:

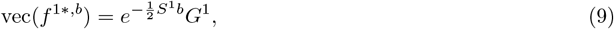

where 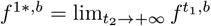.

For the identities represented by the horizontal edges, namely 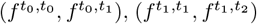, and 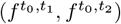, it is not easy to directly formulate and solve the corresponding PDEs. Since we have already established the identities corresponding to the vertical and diagonal edges, we derive those for the horizontal edges indirectly (for details of the derivation, see Supplementary Information A), which yields:

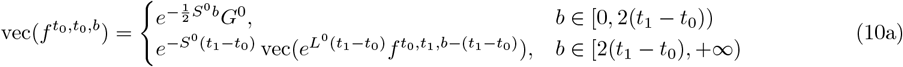

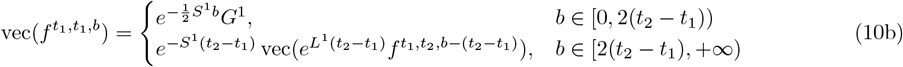

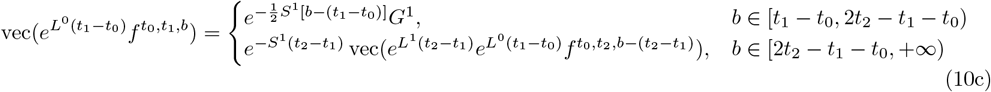

These equations are highly complex both in terms of representation and computation. The difficulty arises from the fact that when we trace lineage 2 in *f* ^*x,y,b*^ along *y*, the process is not purely migrational: we must also account for the location of lineage 1 at the same time point. If lineage 1 happens to be in the same deme as lineage 2, there is a positive probability that they coalesce. Moreover, for lineage 1 to evolve from time *x* to time *y*, it must pass through all changes in the migration rates during the interval [*x, y*]. This is why 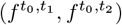, represented by the black horizontal edge in Fig. 2, is particularly intricate, as it depends simultaneously on *L*^0^, *L*^1^, and *γ*^1^.

Since all identities represented by the horizontal edges can be effectively derived from those represented by the vertical and diagonal edges, and are typically much more complicated than the other two categories, it is important to avoid using them for inference. In a demographic inference framework based on pairwise branch lengths, one should rely only on identities represented by the vertical edges when estimating backward migration rates, and use both vertical and diagonal edges for the joint inference of migration rates and coalescence rates.

### Identities relating expected pairwise branch lengths

The identities among the probability density functions (PDFs) of pairwise branch lengths naturally extend to summary statistics derived from them. For example, consider the expected pairwise branch length 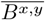. In the structured coalescent framework, the analogous quantity—expected pairwise coalescence times—is directly related to sample covariance structure (McVean, 2009), a connection exploited by methods including EEMS (Petkova et al., 2016), FEEMS (Marcus et al., 2021), and FRAME (Shen and Novembre, 2025). As we demonstrate in Supplementary Information D, such a connection can extend to serially sampled data with slight modifications. Furthermore, advances in tree sequence reconstruction now allow for the direct estimation of 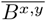 from inferred branch lengths, though this requires careful consideration of the demographic assumptions inherent in the reconstruction process. Here, we establish the identities among expected pairwise branch lengths in the structured serial coalescent.

Similar to the last section, we study a three-epoch toy model. The 8 identities we would like to study are 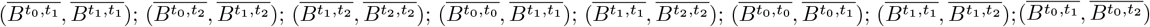. These identities correspond to edges in Fig. 3.

**Fig. 3.**
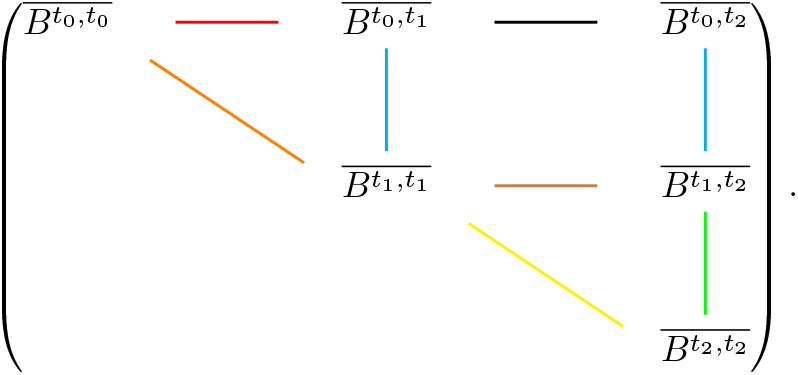
Graphical representation of the identities relating expected pairwise branch lengths.

We now study 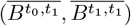. When *x* ∈ [*t*_0_, *t*_1_), this expectation satisfies the partial differential equation:

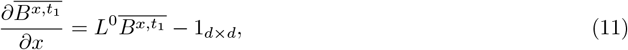

with boundary condition given by 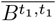. Using the identities *L*1_*d×d*_ = 0 and 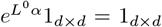 for any *α*, the solution is given by:

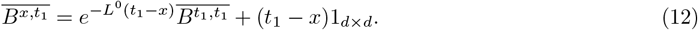

Taking the limit as *x* → *t*_0_, we get 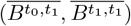. Following the same logic, the identities represented by other vertical edges can also be computed. These identities can be jointly written as:

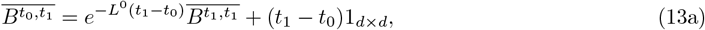

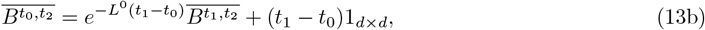

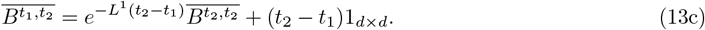

The identities corresponding to the diagonal edges can also be derived directly. Let 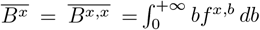 denote the matrix of expected pairwise branch lengths when both lineages are sampled at time *x*. For *x* ∈ [*t*_0_, *t*_1_), this satisfies the ordinary differential equation:

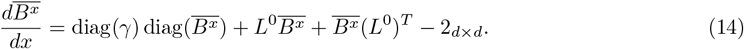

Vectorizing and solving this equation gives:

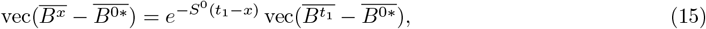

where 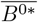 is the equilibrium expected pairwise branch length under the parameter set (*L*^0^, *γ*^0^) and satisfies the equation:

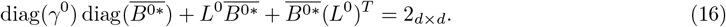

Now letting *x* → *t*_0_ and applying the same logic to the epoch [*t*_1_, *t*_2_), we obtain the joint equation for the identities represented by the diagonal edges:

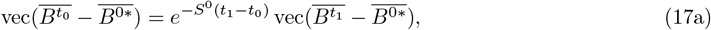

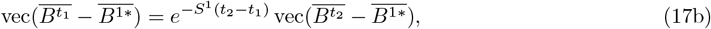

where 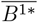 satisfies:

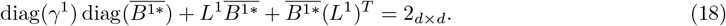

The identities corresponding to the horizontal edges can also be derived indirectly (for a detailed derivation, see Supplementary Information B), and can be written as:

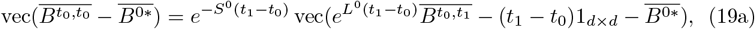

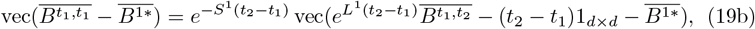

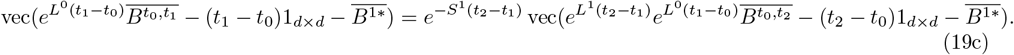

As expected, the identities corresponding to the vertical edges depend only on migration rates and are relatively easy to compute, whereas those corresponding to the horizontal edges are the most complex in terms of both representation and computation.

### Identities relating survival functions of LPSC segment lengths

In the classical coalescent as well as the structured coalescent, it is well known that the length distribution of long pairwise shared coalescence segments (LPSC segments, or identity-by-descent tracts) is determined by the distribution of pairwise coalescence times (Palamara et al., 2012; Carmi et al., 2013) and can thus be used for demographic inference. This is the idea underlying MAPS (Al-Asadi et al., 2019), which infers the migration surfaces and coalescence rates based on the LPSC length distribution. In the structured serial coalescent, following the logic of the previous two sections, we establish the identities relating the survival functions of the LPSC segment lengths.

Let *ρ*^*x,y,µ*^ be the matrix of survival functions such that its (*i, j*)-th element is the probability that an LPSC segment between two randomly sampled lineages from deme *i* at time *x* and deme *j* at time *y* has length larger than *µ*. The probability that a random LPSC segment has length larger than *µ* given the pairwise branch length *b* and recombination rate *r* is simply *e*^*−rbµ*^. Then we obtain:

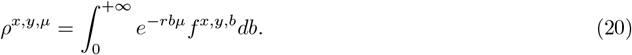

Here again we employ the three-epoch toy model, and we would like to investigate the 8 identities 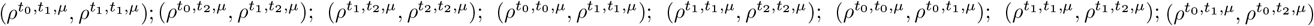. Again, these 8 identities correspond to the 8 edges in Fig. 4.

**Fig. 4.**
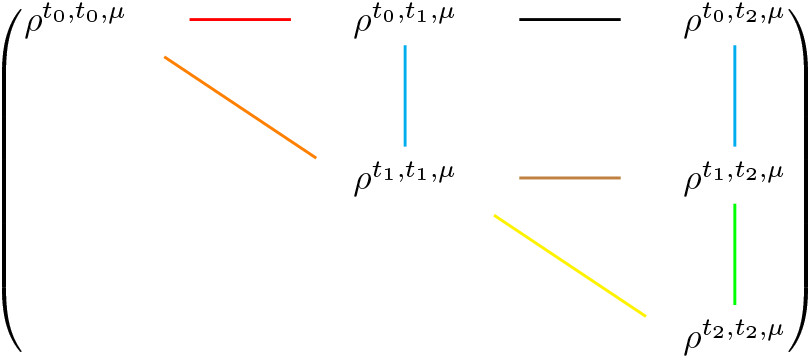
Graphical representation of the identities relating survival functions of LPSC segment lengths.

The easiest way to study the identities represented by the vertical edges here is to write out the integrals and use the identities relating the probability density functions, instead of solving the PDEs. Writing *L*^0^ + *rµI, L*^1^ + *rµI* as 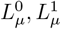 and using Equations (5a), (5b), and (5c) yields:

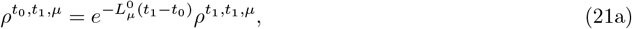

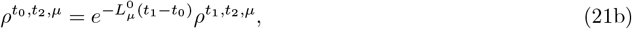

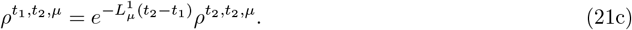

For the identities represented by the diagonal edges, we can also derive analogous expressions. Applying Equations (8a) and (8b), with *ρ*^*x,x,µ*^ = *ρ*^*x,µ*^, 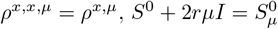, and 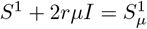, we obtain:

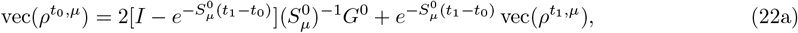

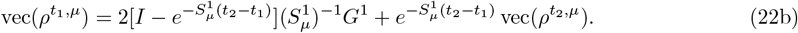

Finally, the identities corresponding to the horizontal edges can also be derived indirectly, which are given by:

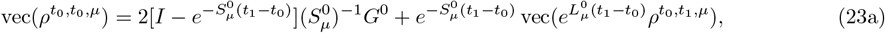

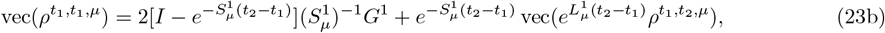

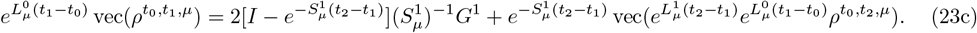

A step-by-step derivation of the identities presented in this section can be found in the Supplementary Information C. Once again, we observe that the identities corresponding to the vertical edges are the most well-behaved, while those corresponding to the horizontal edges are the most complex in terms of both representation and computation.

### Applications for inference

Here we show how the above identities can be used to carry out parameter inference. Similar to the previous sections, we set up a proof-of-concept example by assuming that samples are available at three time points, *t*_0_, *t*_1_, and *t*_2_, and that the goal is to infer the demographic parameters in the time intervals [*t*_0_, *t*_1_) and [*t*_1_, *t*_2_). Both rely on 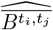, i.e., the estimates obtained for the 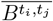 matrices. Such estimates might be based on corresponding data such as inferred branch lengths from reconstructed tree sequences or observed covariance structure in SNP genotypes.

A more fully probabilistic approach that propagates uncertainty through the inference chain would be preferable, but we leave that for future work, and here illustrate how in principle inference of time-varying migration and coalescent rates is possible using the identities derived above in a sequential fashion.

### Pure migration-rate inference

For pure migration-rate inference, we define a simple three-step protocol:

1. Obtain the estimates 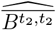 from the corresponding data. This estimation can be straightforward if genetic samples are available from every deme. However, for cases involving vacant demes (those without samples), imputation becomes necessary. One strategy is to use spatial imputation, which leverages geographical proximity to infer values for the vacant demes based on the directly estimated submatrix of 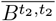 that represents the demes with samples. Alternatively, a model-based imputation approach can be employed. This method operates under the assumption that the stochastic process governing lineages before time *t*_2_ was driven by a set of equilibrium migration rates (*L*^*∞*^) and coalescence rates (*γ*^*∞*^). In other words, it assumes 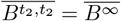, where the equilibrium matrix is defined by the equation:

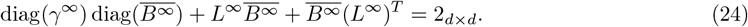 Under this assumption, established equilibrium-based methods, such as FRAME (Shen and Novembre, 2025), can be used to first infer the parameters *L*^*∞*^ and *γ*^*∞*^, which in turn provides an estimate for the complete 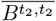 matrix.
2. Estimate 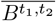 and *L*^1^ based on corresponding data and plugging in 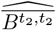 using Equation (13c). From Equation (13c), we know 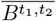is a function of 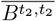and *L*^1^. Since in the first step we already have the inferred 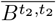, we can write 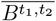solely as a function of *L*^1^. Consequently, a connection between *L*^1^ and the corresponding data is mediated through 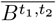. This established identity allows for the estimation of *L*^1^. Notably, if vacant demes prevent the direct estimation of 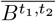from corresponding data, a model-based imputation step can be performed simultaneously with the estimation of *L*^1^.
3. Estimate 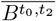and *L*^0^ based on the corresponding data and estimated 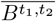using Equation (13b). Here the logic is similar to that in Step 2. Importantly, Equation (13a) can also be used to help estimate *L*^0^ if 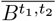can be properly estimated from the corresponding data.

### Joint inference of migration rates and coalescence rates

While the joint inference of migration and coalescence rates involves additional steps, the procedure for estimating migration rates remains consistent with the gene-flow-only protocol. Therefore, for these established steps, we will simply repeat the instructions without further explanation. New procedures specific to joint inference will be explained in detail. The protocol in the proof-of-concept example is as follows:

1. Estimate the 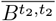 matrix from the corresponding data.
2. Estimate 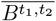and *L*^1^ based on corresponding data and estimated 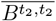using Equation (13c).
3. Estimate 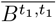and *γ*^1^ based on corresponding data, estimated 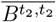, and estimated *L*^1^ using Equation (17b). The idea here again is to use 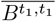as a mediator to connect *γ*^1^ to corresponding data. It is important to mention that we directly use the estimate of *L*^1^ from the previous step instead of estimating it jointly with *γ*^1^.
4. Estimate 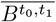, 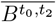, and *L*^0^ based on corresponding data, estimated 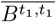, and estimated 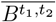 using Equations (13a) and (13b). The underlying logic is analogous to Step 3 of the gene-flow inference protocol. The key distinction is that 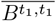is now already estimated from the previous step, which simplifies the direct application of Equation (13a). Again, in practice, researchers can either choose the more convenient formulation or leverage both.
5. Estimate 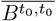 and *γ*^0^ based on the corresponding data and estimated 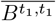using Equation (17a). This step follows the same logic as Step 3.

While the proof-of-concept example demonstrates the core framework, it can be generalized to cases with more than two time slices and other data types, such as sample covariance structures or LPSC segment data.

### Simulation

Here, we test the feasibility of inferring pure migration rates using the inference protocol outlined above. Similar to the previous analysis, we consider a model with three epochs, [*t*_0_, *t*_1_), [*t*_1_, *t*_2_), and [*t*_2_, *t*_3_). In each epoch, we impose a different migration topology and assign random effective population sizes (i.e., coalescence rates). For each combination of topologies, we sample 10 haplotype lineages per deme per epoch and use msprime (Baumdicker et al., 2022) to simulate 10, 000 tree sequences. Parameters are then inferred from the pairwise branch length data of these simulated datasets as follows:

- Obtain 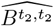 using the method of moments based on the observed branch lengths in the simulated data. For simplicity, in this step we directly use the sample mean of pairwise branch length, i.e., 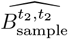, since we have samples from all the demes.
- Estimate 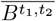and *L*^1^ based on corresponding data and estimated 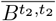using Equation (13c). From the last step, we already have 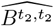, and we first obtain a preliminary estimate of 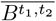by taking the sample mean of the pairwise branch lengths, denoted 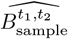. We then infer *L*^1^ by solving an optimization problem with objective 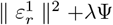, where 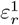 is the relative error matrix of Equation (13c) computed from 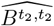 and 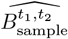, Ψ is a smoothness penalty on edge weights, and *λ* is a hyperparameter selected by cross-validation (see Supplementary Information E for details). Once we obtain 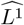, we refine our estimate of 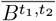using:

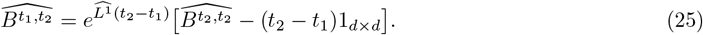
- Estimate 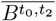and *L*^0^ based on corresponding data and estimated 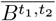using Equation (13b). The procedure follows the same logic as in the last step: we first obtain a direct sample-mean estimate of 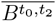, and then infer *L*^0^ by solving an analogous optimization problem. Finally, we refine 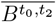 using the resulting 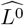.

The results of the simulation study are shown in Fig. 5. Figs. 5(a–c) present the ground-truth topologies for each epoch. In Fig. 5a, the sequence of topologies is as follows: during epoch [*t*_0_, *t*_1_], we have a topology of large-scale directionally migrating lineages, where lineages from the left boundary migrate backward to the right boundary; during epoch [*t*_1_, *t*_2_], the topology is large-scale spatially converging lineages, where lineages converge backward to the center; and during epoch [*t*_2_, + ∞), the topology consists of a mixture of small-scale patterns. We denote these by topology 1, topology 2, and topology 3, respectively. Thus, Fig. 5a corresponds to the topology sequence 1 → 2 → 3. Fig. 5b and Fig. 5c show two other settings obtained by rotating this sequence (2 → 3 → 1 for Fig. 5b and 3 → 1 → 2 for Fig. 5c). The detailed settings and visualization of simulation parameters are provided in Supplementary Information F and Supplementary Figs. 1–3.

**Fig. 5.**
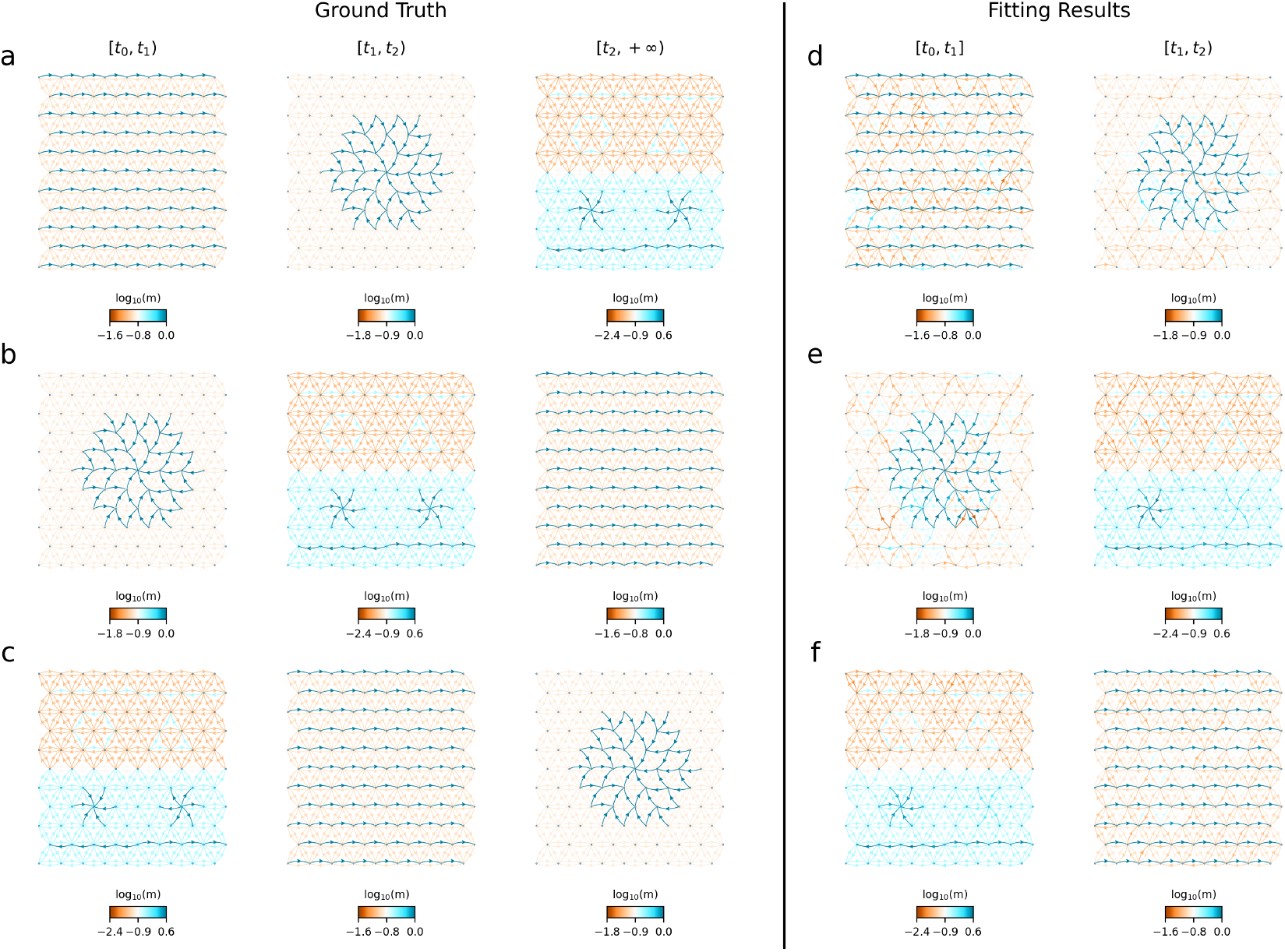
Simulation results. The first row (**a–c**) presents the ground truth topology sequences across three epochs. We define three distinct migration topologies: topology 1 is large-scale directionally migrating lineages, topology 2 is large-scale spatially converging lineages, and topology 3 is a mixture of small scale patterns. Panel (a) shows the sequence 1 → 2 →3, panel (b) the sequence 2 →3 →1, and panel (c) the sequence 3 →1 →2. The second row (**d–f**) shows the corresponding inferred migration patterns for each of the three settings.

Figs. 5(d–f) show the inferred migration patterns. In all three settings, our method largely recovers the underlying structures, including many of the small-scale patterns. The only exception is the spatially diverging lineages in topology 3, which are less clearly recovered. This might be due to the fact that the center of the spatially diverging lineages is visited much less than the center of the spatially converging lineages.

In previous studies such as EEMS (Petkova et al., 2016), FEEMS (Marcus et al., 2021), and FRAME (Shen and Novembre, 2025), migration rates could only be inferred in a relative sense. Here, since we directly use the data of pairwise branch lengths, we can estimate the absolute migration rates.

## Discussion

This study establishes an analytical foundation for pairwise genealogical processes under the structured serial coalescent. By deriving and solving evolution equations for pairwise branch length distributions and their expectations, we characterize fundamental identities governing the interplay of migration and coalescence across time intervals. Our systematic classification identifies distinct classes—graphically represented as edge classes in Figs. 2–4—and evaluates their inference utility through parametric dependencies and computational complexity. This analysis reveals that identities represented by the vertical edges depend exclusively on migration rates, which is also validated in our analysis of identities among length distributions of LPSC segments. This decoupling of spatial dynamics from coalescence processes provides a powerful means of ad-dressing identifiability issues arising from the interaction between migration and coalescence, while enabling scalable inference of migration dynamics.

Leveraging this decoupling, we propose inference frameworks that fully exploit the migration-exclusive nature of identities represented by the vertical edges. Our approach employs a forward-in-time sequential inference strategy: beginning with inferring/imputing the focal quantity related to pairwise branch lengths and parameters at the oldest time point, we sequentially reconstruct migration rates (and coalescence rates) using identities represented by the vertical (and diagonal) edges. The framework naturally accommodates time-stratified sampling and can be adapted to alternative summary statistics.

In addition, since *e*^*L*Δ*t*^ is a stochastic matrix, by replacing it with a probability matrix *P*, the method can also be used to infer the proportion of ancestry from different demes over any discrete time interval Δ*t*. Such an idea has been employed by a recent method (Isacchini et al., 2026) to infer a 9-deme migration network dynamics of ancient Europe. The method the authors used to eliminate genetic drift shares the same philosophy as the ideas emphasized in this paper, since *F*_2_ statistics are based on sample covariance structure, which can be easily translated into expected pairwise coalescence times (Peter, 2016, also see Supplementary Information D).

Despite these advances, several challenges remain for future work. First, the optimization and inference procedure we employ a proof-of-concept example, and a more thorough development of a procedure would be necessary to develop formal probabilistic inference methods based on pairwise branch lengths or observable statistics based on them. The approach outlined here assumed direct estimates of pairwise branch lengths would be the basis of inference, and as such, the migration and coalescence parameter estimation accuracy depends critically on reliable estimates of expected pairwise branch lengths. In empirical applications, pairwise branch lengths are estimated from inferred genealogical trees rather than the true underlying trees (as used in the simulations here). The gap between these two sources requires careful evaluation, since most existing tree inference methods assume panmixia and do not incorporate a formal migration model. In addition, imputation procedures used to address demes with no samples may introduce further errors. In a sequential inference framework based on expected pairwise branch lengths, these errors can propagate, and similar propagation issues can arise in SNP-based inference and LPSC-based inference, although the sources of error can differ. Addressing this problem will be particularly important for inference spanning many epochs. One possible strategy to reduce error propagation is a block-wise sequential approach, in which parameters are inferred jointly across multiple consecutive epochs rather than strictly one epoch at a time. Our focus here has been on understanding the theoretical properties of pairwise branch length statistics and demonstrating the potential of such statistics for inference, however a more formal and developed inference method remains a goal of future work.

Second, our proof-of-concept example does not address the issue of long-range migration. This challenge has already been noted in previous studies (Petkova et al., 2016; Marcus et al., 2021; Shen and Novembre, 2025). Strictly local migration renders the migration-rate matrix sparse, and such sparsity can be leveraged for computational efficiency (Marcus et al., 2021) and parameter identifiability (Lundgren and Ralph, 2019). However, a strictly local network may be insufficient, as real populations may also have experienced gene flow from long-range connections. The difficulty is that these long-range sources are not known *a priori* ; if they were, one could simply include them in the network. The FEEMS paper (Marcus et al., 2021) proposed two strategies to address this issue: a “greedy” algorithm akin to TreeMix (Pickrell and Pritchard, 2012) and a combination of the graphical lasso (Friedman et al., 2008) with graph Laplacian smoothing (Wang et al., 2016). The former has already been explored in FEEMSmix (Shastry et al., 2025), whereas the latter remains to be investigated.

Third, to further accelerate inferences, it is important to adopt efficient algorithms to compute high-dimensional matrix exponentials—particularly for matrices with specific sparse structures. The case of pure migration rate estimation requires exponentiating a *d* × *d* matrix (*e*^*Lt*^), while coalescence rate estimation involves a *d*^2^ × *d*^2^ matrix (*e*^*St*^). While directly computing *e*^*Lt*^ is feasible, directly computing *e*^*St*^ can be computationally expensive for large *d*. In this case, computation and approximation strategies that maximally exploit the structure of *S* can be very helpful.

As the inference challenges outlined earlier are properly addressed, this framework may support powerful applications across evolutionary biology, epidemiology, and conservation. In human evolution, it can augment reconstruction of demographic history from ancient DNA, clarifying migration patterns and their connections to cultural and environmental change. In epidemiological research, it could aid the reconstruction of pathogen transmission dynamics across space and time, facilitating the monitoring of disease spread. In conservation genetics, it may help reveal how gene flow among natural populations shifts through time and how these changes are shaped by environmental pressures such as habitat loss, fragmentation, and climate change. Together, these contributions may provide new avenues to analyze population histories where spatial and temporal dimensions interact—from deep evolutionary timescales to contemporary ecological monitoring.

## Acknowledgements

We would like to thank Ahmed Selim, Sherif Negm, and Egor Lappo for helpful discussions and feedback on the manuscript. The research was supported by funding from NIH NIGMS grant R35-GM149521 to John Novembre.

## Code Availability

The code to reproduce all results is available at https://github.com/ShenHaotv/When-migration-leaves-a-clean-trace.

