## Supplementary Information for "When migration leaves a clean trace: Decoupling migration from coalescence in the structured serial coalescent"

### Contents

**A Identities among probability density functions of pairwise branch length 2**

**B Identities among expected pairwise branch lengths 4**

**C Identities among survival functions of LPSC segment lengths 5**

**D Expected pairwise branch length and sample covariance structure under the structured serial coalescent 7**

**E Objective function and cross validation 9**

**F Simulation settings 11**

**G Supplemental Figures 12**

### A Identities among probability density functions of pairwise branch length

In the main text, we have established the identities corresponding to the vertical edges and diagonal edges, namely $(f^{t_0,t_1}, f^{t_1,t_1})$;$(f^{t_0,t_2}, f^{t_1,t_2})$; $(f^{t_1,t_2}, f^{t_2,t_2})$; $(f^{t_0,t_0}, f^{t_1,t_1})$ and $(f^{t_1,t_1}, f^{t_2,t_2})$. These identities are:

$$f^{t_0,t_1,b} = e^{-L^0(t_1-t_0)} f^{t_1,t_1,b-(t_1-t_0)}, \tag{S1a}$$

$$f^{t_0,t_2,b} = e^{-L^0(t_1-t_0)} f^{t_1,t_2,b-(t_1-t_0)}, \tag{S1b}$$

$$f^{t_1,t_2,b} = e^{-L^1(t_2-t_1)} f^{t_2,t_2,\,b-(t_2-t_1)}; \tag{S1c}$$

$$\mathrm{vec}(f^{t_0,b}) = \begin{cases} e^{-\frac{1}{2}S^0 b} G^0 & b \in [0, 2(t_1-t_0)) \\ e^{-S^0(t_1-t_0)}\,\mathrm{vec}(f^{t_1,b-2(t_1-t_0)}) & b \in [2(t_1-t_0), +\infty) \end{cases} \tag{S2a}$$

$$\mathrm{vec}(f^{t_1,b}) = \begin{cases} e^{-\frac{1}{2}S^1 b} G^1 & b \in [0, 2(t_2-t_1)) \\ e^{-S^1(t_2-t_1)}\,\mathrm{vec}(f^{t_2,b-2(t_2-t_1)}) & b \in [2(t_2-t_1), +\infty) \end{cases} \tag{S2b}$$

We now derive the expressions for the identities corresponding to the horizontal edges, namely $(f^{t_0,t_0}, f^{t_0,t_1})$, $(f^{t_1,t_1}, f^{t_1,t_2})$, and $(f^{t_0,t_1}, f^{t_0,t_2})$. We begin with $(f^{t_0,t_0}, f^{t_0,t_1})$. From equation (S1a), we have

$$\mathrm{vec}(f^{t_0,t_1,b}) = \mathrm{vec}(e^{-L^0(t_1-t_0)} f^{t_1,t_1,b-(t_1-t_0)}). \tag{S3}$$

Combining this expression with (S2a), and performing the reparameterization, we obtain the expression for $(f^{t_0,t_0}, f^{t_0,t_1})$:

$$\mathrm{vec}(f^{t_0,t_0,b}) = \begin{cases} e^{-\frac{1}{2}S^0 b} G^0 & b \in [0, 2(t_1-t_0)) \\ e^{-S^0(t_1-t_0)}\,\mathrm{vec}(e^{L^0(t_1-t_0)} f^{t_0,t_1,b-(t_1-t_0)}) & b \in [2(t_1-t_0), +\infty) \end{cases} \tag{S4}$$

Similarly, the identity $(f^{t_1,t_1}, f^{t_1,t_2})$ can be expressed as

$$\mathrm{vec}(f^{t_1,t_1,b}) = \begin{cases} e^{-\frac{1}{2}S^1 b} G^1 & b \in [0, 2(t_2-t_1)) \\ e^{-S^1(t_2-t_1)}\,\mathrm{vec}(e^{L^1(t_2-t_1)} f^{t_1,t_2,b-(t_2-t_1)}) & b \in [2(t_2-t_1), +\infty) \end{cases} \tag{S5}$$

From equation (S1b), we also have

$$\mathrm{vec}(f^{t_0,t_2,b}) = \mathrm{vec}(e^{-L^0(t_1-t_0)} f^{t_1,t_2,b-(t_1-t_0)}). \tag{S6}$$

Combining the last two equations with equation (S1a) and after reparameterization we obtain the expression for $(f^{t_0,t_1}, f^{t_0,t_2})$:

$$\begin{aligned} &\operatorname{vec}(e^{L^0(t_1-t_0)} f^{t_0,t_1,b}) \\ =&\begin{cases} e^{-\frac{1}{2}S^1[b-(t_1-t_0)]}G^1 & b \in [t_1-t_0, 2t_2-t_1-t_0) \\ e^{-S^1(t_2-t_1)} \operatorname{vec}(e^{L^1(t_2-t_1)} e^{L^0(t_1-t_0)} f^{t_0,t_2,b-(t_2-t_1)}) & b \in [2t_2-t_1-t_0, +\infty) \end{cases} \end{aligned} \tag{S7}$$

#### B Identities among expected pairwise branch lengths

In the main text, we have established the identities corresponding to the vertical edges and diagonal edges, namely $(\overline{B^{t_0,t_1}}, \overline{B^{t_1,t_1}})$; $(\overline{B^{t_0,t_2}}, \overline{B^{t_1,t_2}})$; $(\overline{B^{t_1,t_2}}, \overline{B^{t_2,t_2}})$; $(\overline{B^{t_0,t_0}}, \overline{B^{t_1,t_1}})$ and $(\overline{B^{t_1,t_1}}, \overline{B^{t_2,t_2}})$:

$$\overline{B^{t_0,t_1}} = e^{-L^0(t_1-t_0)}\overline{B^{t_1,t_1}} + (t_1 - t_0)1_{d\times d}, \tag{S8a}$$

$$\overline{B^{t_0,t_2}} = e^{-L^0(t_1-t_0)}\overline{B^{t_1,t_2}} + (t_1 - t_0)1_{d\times d}, \tag{S8b}$$

$$\overline{B^{t_1,t_2}} = e^{-L^1(t_2-t_1)}\overline{B^{t_2,t_2}} + (t_2 - t_1)1_{d\times d}; \tag{S8c}$$

$$\operatorname{vec}(\overline{B^{t_0}} - \overline{B^{0*}}) = e^{-S^0(t_1-t_0)} \operatorname{vec}(\overline{B^{t_1}} - \overline{B^{0*}}), \tag{S9a}$$

$$\operatorname{vec}(\overline{B^{t_1}} - \overline{B^{1*}}) = e^{-S^1(t_2-t_1)} \operatorname{vec}(\overline{B^{t_2}} - \overline{B^{1*}}). \tag{S9b}$$

Here we derive the expression for $(\overline{B^{t_0,t_0}}, \overline{B^{t_0,t_1}})$; $(\overline{B^{t_1,t_1}}, \overline{B^{t_1,t_2}})$; $(\overline{B^{t_0,t_1}}, \overline{B^{t_0,t_2}})$, which correspond to the horizontal edges. We first study $(\overline{B^{t_0,t_0}}, \overline{B^{t_0,t_1}})$. From equation (S8a), we have

$$\operatorname{vec}(\overline{B^{t_0,t_1}}) = \operatorname{vec}(e^{-L^0(t_1-t_0)}\overline{B^{t_1,t_1}} + (t_1 - t_0)1_{d\times d}), \tag{S10}$$

and from equation (S9a),

$$\operatorname{vec}(\overline{B^{t_0,t_0}} - \overline{B^{0*}}) = e^{-S^0(t_1-t_0)} \operatorname{vec}(\overline{B^{t_1,t_1}} - \overline{B^{0*}}). \tag{S11}$$

Combining the two gives the expression for $(\overline{B^{t_0,t_0}}, \overline{B^{t_0,t_1}})$:

$$\operatorname{vec}(\overline{B^{t_0,t_0}} - \overline{B^{0*}}) = e^{-S^0(t_1-t_0)} \operatorname{vec}(e^{L^0(t_1-t_0)}\overline{B^{t_0,t_1}} - (t_1-t_0)1_{d\times d} - \overline{B^{0*}}). \tag{S12}$$

In a similar way, the identity $(\overline{B^{t_1,t_1}}, \overline{B^{t_1,t_2}})$ can be written as

$$\operatorname{vec}(\overline{B^{t_1,t_1}} - \overline{B^{1*}}) = e^{-S^1(t_2-t_1)} \operatorname{vec}(e^{L^1(t_2-t_1)}\overline{B^{t_1,t_2}} - (t_2-t_1)1_{d\times d} - \overline{B^{1*}}). \tag{S13}$$

Finally, from equation (S8b), we obtain

$$\operatorname{vec}(\overline{B^{t_0,t_2}}) = \operatorname{vec}(e^{-L^0(t_1-t_0)}\overline{B^{t_1,t_2}} + (t_1 - t_0)1_{d\times d}). \tag{S14}$$

Combining this with equations (S11) and (S13), we derive the expression for $(\overline{B^{t_0,t_1}}, \overline{B^{t_0,t_2}})$:

$$\begin{aligned} &\operatorname{vec}(e^{L^0(t_1-t_0)}\overline{B^{t_0,t_1}} - (t_1 - t_0)1_{d\times d} - \overline{B^{1*}}) \\ =&e^{-S^1(t_2-t_1)} \operatorname{vec}(e^{L^1(t_2-t_1)}e^{L^0(t_1-t_0)}\overline{B^{t_0,t_2}} - (t_2 - t_0)1_{d\times d} - \overline{B^{1*}}). \end{aligned} \tag{S15}$$

### C Identities among survival functions of LPSC segment lengths

Here we give a detailed derivation of identities among survival functions of LPSC segment lengths given in the main text. From main text we have

$$\rho^{x,y,\mu} = \int_0^{+\infty} e^{-rb\mu} f^{x,y,b} db. \tag{S16}$$

Writing $L^0 + r\mu I, L^1 + r\mu I$ as $L^0_\mu, L^1_\mu$ and using equation (S1a) yields

$$\begin{aligned}\rho^{t_0,t_1,\mu} &= \int_0^{+\infty} e^{-rb\mu} f^{t_0,t_1,b} db \\ &= e^{-L^0(t_1-t_0)} \int_0^{+\infty} e^{-rb\mu} f^{t_1,t_1,b-(t_1-t_0)} db \\ &= e^{-L^0_\mu(t_1-t_0)} \rho^{t_1,t_1,\mu}\end{aligned} \tag{S17}$$

Applying equations (S1b) and (S1c) similarly gives

$$\rho^{t_0,t_2,\mu} = e^{-L^0_\mu(t_1-t_0)} \rho^{t_1,t_2,\mu} \tag{S18}$$

and

$$\rho^{t_1,t_2,\mu} = e^{-L^1_\mu(t_2-t_1)} \rho^{t_2,t_2,\mu}. \tag{S19}$$

The three equations above show the identities represented by the vertical edges, namely $(\rho^{t_0,t_1,\mu}, \rho^{t_1,t_1,\mu})$, $(\rho^{t_0,t_2,\mu}, \rho^{t_1,t_2,\mu})$ and $(\rho^{t_1,t_2,\mu}, \rho^{t_2,t_2,\mu})$. We can perform a similar analysis for the identities represented by the diagonal edges, i.e. $(\rho^{t_0,t_0,\mu}, \rho^{t_1,t_1,\mu})$ and $(\rho^{t_1,t_1,\mu}, \rho^{t_2,t_2,\mu})$. Let $\rho^{x,x,\mu} = \rho^{x,\mu}$ and $S^0 + 2r\mu I = S^0_\mu$ . Using equation (S2a), we obtain

$$\begin{aligned}\operatorname{vec}(\rho^{t_0,\mu}) &= \int_0^{2(t_1-t_0)} e^{-rb\mu} e^{-\frac{1}{2}S^0 b} G^0 db \\ &+ \int_{2(t_1-t_0)}^{+\infty} e^{-rb\mu} e^{-S^0(t_1-t_0)} \operatorname{vec}(f^{t_1,b-2(t_1-t_0)}) db \\ &= 2[I - e^{-S^0_\mu(t_1-t_0)}](S^0_\mu)^{-1} G^0 + e^{-S^0_\mu(t_1-t_0)} \operatorname{vec}(\rho^{t_1,\mu}).\end{aligned} \tag{S20}$$

Similarly, let $S^1 + 2r\mu I = S^1_\mu$, and using equation (S2b) we have

$$\operatorname{vec}(\rho^{t_1,\mu}) = 2[I - e^{-S^1_\mu(t_2-t_1)}](S^1_\mu)^{-1} G^1 + e^{-S^1_\mu(t_2-t_1)} \operatorname{vec}(\rho^{t_2,\mu}). \tag{S21}$$

To obtain the expression for the identity $(\rho^{t_0,t_0,\mu}, \rho^{t_0,t_1,\mu})$, we rewrite equation (S20) as

$$\operatorname{vec}(\rho^{t_0,t_0,\mu}) = 2[I - e^{-S^0_\mu(t_1-t_0)}](S^0_\mu)^{-1} G^0 + e^{-S^0_\mu(t_1-t_0)} \operatorname{vec}(\rho^{t_1,t_1,\mu}). \tag{S22}$$

Combining this equation with equation (S17), we obtain

$$\mathrm{vec}(\rho^{t_0,t_0,\mu}) = 2[I - e^{-S^0_\mu(t_1-t_0)}](S^0_\mu)^{-1}G^0 + e^{-S^0_\mu(t_1-t_0)}\,\mathrm{vec}(e^{L^0_\mu(t_1-t_0)}\rho^{t_0,t_1,\mu}). \tag{S23}$$

Similarly, the identity $(\rho^{t_1,t_1,\mu}, \rho^{t_1,t_2,\mu})$ can be expressed as

$$\mathrm{vec}(\rho^{t_1,t_1,\mu}) = 2[I - e^{-S^1_\mu(t_2-t_1)}](S^1_\mu)^{-1}G^1 + e^{-S^1_\mu(t_2-t_1)}\,\mathrm{vec}(e^{L^1_\mu(t_2-t_1)}\rho^{t_1,t_2,\mu}). \tag{S24}$$

Combining equation (S24) with equation (S17) and (S18), we obtain the expression for $(\rho^{t_0,t_1,\mu}, \rho^{t_0,t_2,\mu})$:

$$\begin{aligned} & e^{L^0_\mu(t_1-t_0)}\,\mathrm{vec}(\rho^{t_0,t_1,\mu}) \\ =& 2[I - e^{-S^1_\mu(t_2-t_1)}](S^1_\mu)^{-1}G^1 + e^{-S^1_\mu(t_2-t_1)}\,\mathrm{vec}(e^{L^1_\mu(t_2-t_1)}e^{L^0_\mu(t_1-t_0)}\rho^{t_0,t_2,\mu}) \end{aligned} \tag{S25}$$

### D Expected pairwise branch length and sample covariance structure under the structured serial coalescent

We establish the relationship between the expected pairwise branch lengths and sample covariance structure under the structured serial coalescent, following the framework of McVean (2009). For a given SNP, let the allele states of two haploid samples $X$ and $Y$ be $Z_X, Z_Y \in \{0, 1\}$. Conditioned on the event $S$ that exactly one mutation occurs in the genealogy, and taking the limit as the mutation rate $\theta \to 0$, the expectation of $Z_X$ can be expressed as

$$E(Z_X|S) = \frac{\Pr(Z_X = 1, S)}{P(S)} = \lim_{\theta \to 0} \frac{E(\theta(t_G^{MRCA} - t_X)e^{-\theta B_{tot}})}{E(\theta B_{tot} e^{-\theta B_{tot}})} = \frac{\overline{t_G^{MRCA}} - t_X}{\overline{B_{tot}}}, \tag{S26}$$

where $\overline{t_G^{MRCA}}$ is the expected time of the most recent common ancestor of the genealogy, and $\overline{B_{tot}}$ is the expected total branch length. Similarly, the second moments are given by

$$E(Z_Y|S) = \frac{\overline{t_G^{MRCA}} - t_Y}{\overline{B_{tot}}}, \tag{S27a}$$

$$E(Z_X Z_Y|S) = \frac{\overline{t_G^{MRCA}} - \overline{t_{XY}^{MRCA}}}{\overline{B_{tot}}} = \frac{2\overline{t_G^{MRCA}} - (t_X + t_Y) - \overline{B_{XY}}}{2\overline{B_{tot}}}, \tag{S27b}$$

where $\overline{B_{XY}}$ is the expected pairwise branch length between $X$ and $Y$. Since the sampling times $t_X, t_Y$ are known parameters, the connection between $E(Z_X Z_Y|S)$ and $\overline{B_{XY}}$ is established by treating $\overline{B_{tot}}$ as a nuisance parameter and eliminating $\overline{t_G^{MRCA}}$ through contrasts, as practiced in EEMS (Petkova et al., 2016), FEEMS (Marcus et al., 2021), and FRAME (Shen and Novembre, 2025).

This formulation is also closely related to the $F_2$ statistics. For two demes $i$ and $j$ sampled at times $t_1, t_2$ with allele frequencies $f_i^{t_1}, f_j^{t_2}$ and sample sizes $n_i^{t_1}, n_j^{t_2}$, extending from the logic above, we have:

$$E(f_i^{t_1} f_j^{t_2}|S) = \frac{2\overline{t_G^{MRCA}} - (t_1 + t_2) - (1 - \frac{\delta_{ij}\delta_{t_1 t_2}}{n_i^{t_1}})\overline{B_{ij}^{t_1,t_2}}}{2\overline{B_{tot}}}. \tag{S28}$$

where $\delta_{xy} = 1$ if $x = y$, and 0 otherwise.

Under the large sample size approximation ($n \to \infty$), the expected squared difference becomes

$$E((f_i^{t_1} - f_j^{t_2})^2|S) \approx \frac{2\overline{B_{ij}^{t_1,t_2}} - \overline{B_{ii}^{t_1,t_1}} - \overline{B_{jj}^{t_2,t_2}}}{2\overline{B_{tot}}}. \tag{S29}$$

Now assume $t_1 < t_2$ (backward in time), and let $F^{t_1,t_2}$ be the matrix where each entry is $F_{ij}^{t_1,t_2} = E((f_i^{t_1} - f_j^{t_2})^2|S)$. Using the equation derived in the main text

$$\overline{B^{t_1,t_2}} = e^{-L^1(t_2-t_1)}\overline{B^{t_2,t_2}} + (t_2-t_1)1_{d\times d}, \tag{S30}$$

we can express the components of $F^{t_1,t_2}$ as

$$F_{ij}^{t_1,t_2} \approx \frac{2[e^{-L^1(t_2-t_1)}\overline{B^{t_2,t_2}}]_{ij} + 2(t_2-t_1) - \overline{B_{ii}^{t_1,t_1}} - \overline{B_{jj}^{t_2,t_2}}}{\overline{2B_{tot}}}, \tag{S31a}$$

$$F_{ii}^{t_1,t_2} \approx \frac{2[e^{-L^1(t_2-t_1)}\overline{B^{t_2,t_2}}]_{ii} + 2(t_2-t_1) - \overline{B_{ii}^{t_1,t_1}} - \overline{B_{ii}^{t_2,t_2}}}{\overline{2B_{tot}}}. \tag{S31b}$$

Taking the difference $F_{ij}^{t_1,t_2} - F_{ii}^{t_1,t_2}$ eliminates the terms involving $t_1$ and the temporal offset $(t_2 - t_1)$:

$$F_{ij}^{t_1,t_2} - F_{ii}^{t_1,t_2} \approx \frac{2[e^{-L^1(t_2-t_1)}\overline{B^{t_2,t_2}}]_{ij} - 2[e^{-L^1(t_2-t_1)}\overline{B^{t_2,t_2}}]_{ii} + \overline{B_{ii}^{t_2,t_2}} - \overline{B_{jj}^{t_2,t_2}}}{\overline{2B_{tot}}}. \tag{S32}$$

For samples at $t_2$, the matrix $F^{t_2,t_2}$ is given by:

$$F^{t_2,t_2} \approx \frac{1}{\overline{2B_{tot}}}(2\overline{B^{t_2,t_2}} - \mathrm{diag}\{\overline{B^{t_2,t_2}}\}1_{d\times d} - 1_{d\times d}\,\mathrm{diag}\{\overline{B^{t_2,t_2}}\}). \tag{S33}$$

By applying the migration operator $e^{-L^1(t_2-t_1)}$ to $F^{t_2,t_2}$ and calculating the row-wise difference, we obtain:

$$\begin{aligned} &[e^{-L^1(t_2-t_1)}F^{t_2,t_2}]_{ij} - [e^{-L^1(t_2-t_1)}F^{t_2,t_2}]_{ii} \\ &\approx \frac{2[e^{-L^1(t_2-t_1)}\overline{B^{t_2,t_2}}]_{ij} - 2[e^{-L^1(t_2-t_1)}\overline{B^{t_2,t_2}}]_{ii} + \overline{B_{ii}^{t_2,t_2}} - \overline{B_{jj}^{t_2,t_2}}}{\overline{2B_{tot}}} \\ &\approx F_{ij}^{t_1,t_2} - F_{ii}^{t_1,t_2}. \end{aligned} \tag{S34}$$

We recognize equation S34 as the core formulation used for migration inference in Isacchini et al. (2026), demonstrating that their approach is equivalent to the $F$-statistics representation of our proposed theory.

### E Objective function and cross validation

Here we provide a detailed introduction to the optimization procedure used in the simulation section of the main text.

The lineage migration process in each epoch can be represented by a directed weighted graph. We number the nodes from 1 to $d$, and use the ordered pair $(i, j)$ to represent an edge from node $i$ to node $j$. Let $\Omega$ denote the edge set under this representation, and define

$$\Omega_i = \{(i,t) \in \Omega\} \cup \{(s,i) \in \Omega\},$$

the set of edges connected to node $i$. We assume that $\Omega$ and $\Omega_i$ remain fixed across epochs, while only the edge weights vary.

Suppose we have $\widehat{B^{t_1,t_2}_{\text{sample}}}$ and $\widehat{B^{t_2,t_2}}$ available, and would like to infer $L^1$ using the equation

$$\overline{B^{t_1,t_2}} - (t_2 - t_1)\, 1_{d\times d} = e^{-L^1(t_2-t_1)}\overline{B^{t_2,t_2}}. \tag{S35}$$

In the main text, we assume that samples are available from all demes. For empirical datasets, however, some demes may lack samples, and the same issue also arises in cross-validation. To handle this, suppose we observe $o^1$ demes at time $t_1$ with indices $O^1 = \{\alpha^1_1, \dots, \alpha^1_{o^1}\}$, and $o^2$ demes at time $t_2$ with indices $O^2 = \{\alpha^2_1, \dots, \alpha^2_{o^2}\}$. Then $\widehat{B^{t_1,t_2}_{\text{sample}}}$ is an $o^1 \times o^2$ matrix.

We define selection matrices $C^1 \in \mathbb{R}^{o^1\times d}$ and $C^2 \in \mathbb{R}^{o^2\times d}$, where $C^1_{ij} = 1$ if $j = \alpha^1_i$ (and 0 otherwise), and $C^2_{ij} = 1$ if $j = \alpha^2_i$ (and 0 otherwise). Equation (S35) then becomes

$$C^1\left[\overline{B^{t_1,t_2}} - (t_2 - t_1)\, 1_{d\times d}\right](C^2)^T = C^1\left[e^{-L^1(t_2-t_1)}\overline{B^{t_2,t_2}}\right](C^2)^T. \tag{S36}$$

The data we have at hand corresponding to the left-hand side of the equation is $\widehat{B^{t_1,t_2}_{\text{sample}}} - (t_2 - t_1)1_{o^1\times o^2}$ , which we denote by $A_1$; The data corresponding to the right-hand side of the equation is $C^1[e^{-L^1(t_2-t_1)}\widehat{B^{t_2,t_2}}](C^2)^T$, which we denote by $A_2$. We can then define a matrix of relative error $\varepsilon^1_r \in R^{o^1\times o^2}$, such that

$$(\varepsilon^1_r)_{ij} = \frac{|(A_2 - A_1)_{ij}|}{|(A_1)_{ij}|}. \tag{S37}$$

The optimization problem can then be written as

$$\widehat{L^1} = \underset{L^1\in\mathcal{D}}{\operatorname{argmin}}\,(\|\, \varepsilon^1_r \,\|^2 + \lambda\Psi), \tag{S38}$$

where $\mathcal{D}$ is the region that all edges have positive weight; $\Psi$ is a smoothness penalty defined by

$$\Psi = \frac{1}{2}\sum_{i=1}^{d} \sum_{\substack{(i_1,j_1)\in\Omega_i,\\ (i_2,j_2)\in\Omega_i}} \frac{1}{|\Omega_i|^2}[\log(m_{i_1j_1}) - \log(m_{i_2j_2})]^2, \tag{S39}$$

and $\lambda$ is a hyperparameter controlling the strength of regularization.

For each fixed $\lambda$, we solve the optimization problem using L-BFGS (Byrd et al., 1995), with gradients computed automatically via PyTorch (Paszke et al., 2019). The optimal $\lambda$ is then selected through cross-validation.

Specifically, for a chosen $k$, we partition the observed demes at time $t_1$ into $k$ folds, $O^1_1, \ldots, O^1_k$, and the observed demes at $t_2$ into $O^2_1, \ldots, O^2_k$. For each $i$, we hold out $(O^1_i, O^2_i)$ and train on the remaining demes. Let $\widehat{L^1_{\text{train}}}$ be the estimate obtained from the training set. In this case, only entries of equation (S35) with row indices in $O^1_{\text{train}} = O^1 \setminus O^1_i$ and column indices in $O^2_{\text{train}} = O^2 \setminus O^2_i$ are used for inference. The held-out entries—those with row indices in $O^1_i$ or column indices in $O^2_i$—serve as the test set.

The validation error for fold $i$ is then defined as the sum of squared relative errors of these entries. Or in an equivalent format, let $(\varepsilon^1_r)_{\text{train}}$ be a matrix computed from $A_1 = \widehat{B^{t_1,t_2}_{\text{train}}} - (t_2 - t_1) 1_{|O^1_{\text{train}}| \times |O^2_{\text{train}}|}$ and $A_2 = C^1_{\text{train}}[e^{-L^1(t_2-t_1)}\widehat{B^{t_2,t_2}}](C^2_{\text{train}})^T$, where $\widehat{B^{t_1,t_2}_{\text{train}}}, C^1_{\text{train}}, C^2_{\text{train}}$ represent the expected pairwise branch length matrix and selection matrix for training demes, the validation error can be defined as $\| \varepsilon^1_r \|^2 - \| (\varepsilon^1_r)_{\text{train}} \|^2$.

### F Simulation settings

We use `msprime` (Baumdicker et al., 2022) to simulate the structured serial coalescent process. We define three time points, $t_0 = 0$, $t_1 = 1$, and $t_2 = 2$, which partition the process into three epochs: $[t_0, t_1)$, $[t_1, t_2)$, and $[t_2, t_\infty)$. In each epoch, one of three graph topologies is applied: large-scale directionally migrating lineages, large-scale spatially converging lineages, or a mixture of small-scale patterns (as shown in Fig. 5 of the main text).

In both the large-scale directionally migrating lineages and large-scale spatially converging lineages scenarios, the base migration rate (corresponding to the orange edges) is set to 0.1, while the blue edges have migration rates 1, which are ten times higher. In the topology combining small-scale patterns, the migration rates along the middle row are 0.1. The upper half of the graph has a base migration rate of 0.03, shown as orange edges. The lower half of the graph has a base migration rate of 0.3, shown as blue edges. All patterns exhibit migration rates that are 10 times higher than the surrounding base migration rates. Specifically, the patterns in the upper half of the graph will have a migration rate of 0.3, while patterns in the lower half will have migration rates of 3.

Effective population sizes are chosen at random by sampling log(Ne) uniformly from the interval $[-1, 1]$. The resulting values used in the simulations are visualized in Supplementary Figs. 1–3.

From each deme, 10 lineages are sampled, and the structured serial coalescent simulation is performed with the specified migration and coalescence rates in each epoch. Each simulation yields a tree sequence. We repeat this procedure 10000 times and conduct inference using the pairwise branch lengths computed from the resulting tree sequences.

### G Supplemental Figures

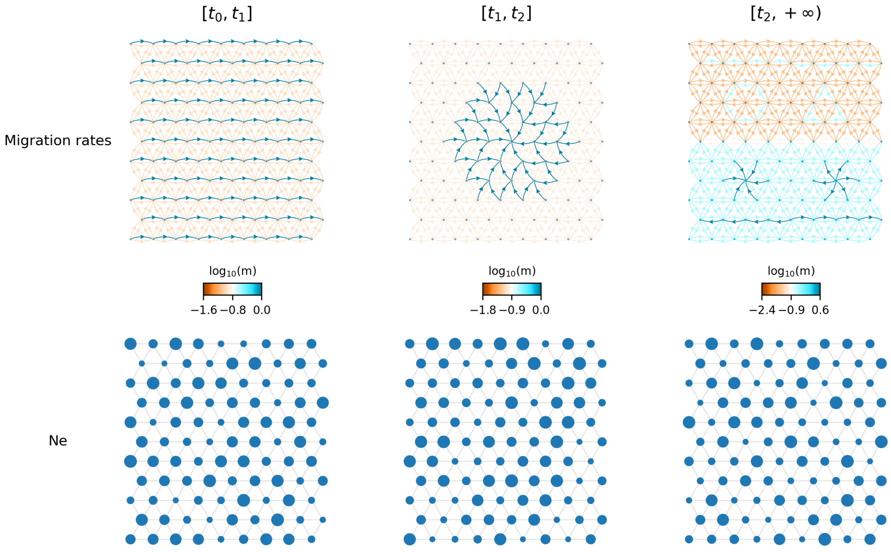


**Fig. S1.** Migration rates and effective population size settings for the topology sequence in Fig. 5a of the main text.

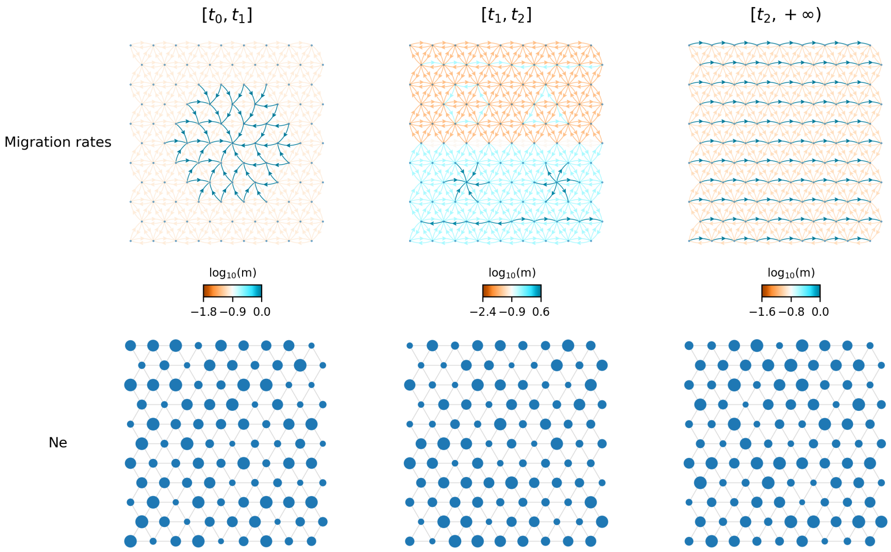


**Fig. S2.** Migration rates and effective population size settings for the topology sequence in Fig. 5b of the main text.

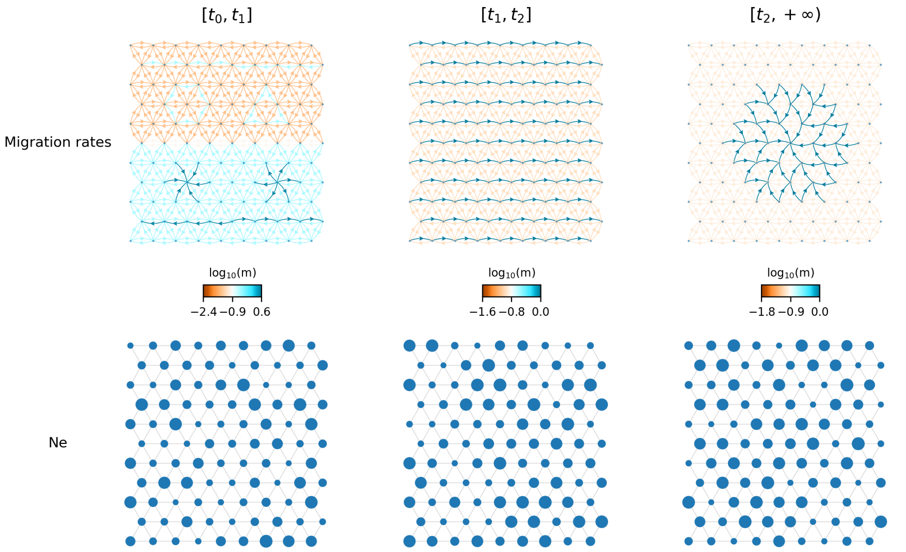


**Fig. S3.** Migration rates and effective population size settings for the topology sequence in Fig. 5c of the main text.
